# HSB-1 inhibition and HSF-1 overexpression trigger overlapping transcriptional changes to promote longevity in *Caenorhabditis elegans*

**DOI:** 10.1101/535864

**Authors:** Surojit Sural, Tzu-Chiao Lu, Seung Ah Jung, Ao-Lin Hsu

**Affiliations:** Department of Molecular and Integrative Physiology, University of Michigan Medical School, Ann Arbor, MI 48109, USA; Research Center for Healthy Aging and Institute of New Drug Development, China Medical University, Taichung, 404, Taiwan; College of Literature, Science, and the Arts, University of Michigan, Ann Arbor, MI 48109, USA; Department of Internal Medicine, Division of Geriatric and Palliative Medicine, University of Michigan Medical School, Ann Arbor, MI 48109, USA; Institute of Biochemistry and Molecular Biology, National Yang Ming University, Taipei, 112, Taiwan

**Keywords:** *C. elegans*, heat shock factor, HSB-1, life span, RNA-Seq

## Abstract

Heat shock factor 1 (HSF-1) is a component of the heat shock response pathway that is induced by cytoplasmic proteotoxic stress. In addition to its role in stress response, HSF-1 also acts as a key regulator of the rate of organismal aging. Overexpression of HSF-1 promotes longevity in *C*. *elegans* via mechanisms that remain less understood. Moreover, genetic ablation of a negative regulator of HSF-1, termed as heat shock factor binding protein 1 (HSB-1), results in *hsf-1*-dependent life span extension in animals. Here we show that in the absence of HSB-1, HSF-1 acquires increased DNA binding activity to its genomic target sequence. Using RNA-Seq to compare the gene expression profiles of the *hsb-1* mutant and *hsf-1* overexpression strains, we found that while more than 1,500 transcripts show ≥1.5-fold upregulation due to HSF-1 overexpression, HSB-1 inhibition alters the expression of less than 500 genes in *C*. *elegans*. Roughly half of the differentially regulated transcripts in the *hsb-1* mutant have altered expression also in *hsf-1* overexpressing animals, with a strongly correlated fold-expression pattern between the two strains. In addition, genes that are upregulated via both HSB-1 inhibition and HSF-1 overexpression include numerous DAF-16 targets that have known functions in longevity regulation. This study identifies how HSB-1 acts as a specific regulator of the transactivation potential of HSF-1 in non-stressed conditions, thus providing a detailed understanding of the role of HSB-1/HSF-1 signaling pathway in transcriptional regulation and longevity in *C*. *elegans*.

## INTRODUCTION

Genes that are involved in stress response pathways have often been implicated in the regulation of organismal longevity in non-stressed conditions (Rodriguez *et al*. 2013; Epel and Lithgow 2014). Comparative studies have shown that cellular resistance to stress is strongly correlated with maximum life span in biologically-related species (Kapahi *et al*. 1999; Harper *et al*. 2007, 2011). Heat shock response (HSR) is one such evolutionarily conserved pathway that is activated in response of various stress conditions such as heat, oxidative damage, proteotoxic insults and bacterial infections (Morimoto 2011). In harsh environmental conditions, HSR triggers the activation of members of the heat shock factor (HSF) family of transcription factors in animals (Akerfelt *et al*. 2010; Morimoto 2011). In vertebrates, the HSF family has four members, namely HSF1–4, while yeast, *C*. *elegans* and *Drosophila* have a sole ortholog of HSF1 (Takii and Fujimoto 2016). In the presence of stress stimuli, the HSF-1 protein acquires post-translational modifications (PTMs), undergoes oligomerization, translocates to the nucleus and shows increased binding to its target sequences in the genome termed as heat shock elements (HSE) (Sarge *et al*. 1993; Chiang *et al*. 2012). Increased HSF-1 activity induces transcriptional upregulation of members of the heat shock protein (HSP) family, which function as molecular chaperones to assist in the folding of nascent polypeptides and prevent the toxic aggregation of misfolded cytosolic proteins (Richter *et al*. 2010). Hence, HSF-1-mediated transcriptional changes influence the survival of organisms in harsh environmental conditions via ameliorating the stress-induced loss of protein homeostasis (McMillan *et al*. 1998; Hsu *et al*. 2003).

HSF-1 has also been found to be a major determinant of organismal life span in non-stressed physiological conditions. *hsf-1* is required for life span extension associated with several longevity-regulating mechanisms, such as insulin/IGF-1-like signaling, target of rapamycin (TOR) signaling and food deprivation (Hsu *et al*. 2003; Morley and Morimoto 2004; Steinkraus *et al*. 2008; Seo *et al*. 2013). In the nematode worm *C*. *elegans*, overexpression of *hsf-1* is sufficient to extend life span and slow the age-related progression of protein aggregation disorders, while RNAi-mediated knockdown of *hsf-1* has the opposite effects on these phenotypes (Hsu *et al*. 2003; Morley and Morimoto 2004). In addition, increased expression of HSF-1 target genes has been shown to be sufficient for extension of life span in *C*. *elegans* and *Drosophila* in non-stressed conditions (Tatar *et al*. 1997; Walker and Lithgow 2003). Initial studies reported that increased survival associated with *hsf-1* overexpression is at least partially due to transcriptional upregulation of small *hsp* genes (Hsu *et al*. 2003). However, a recent study showed that overexpression of a modified form of HSF-1 extended life span of animals without affecting their ability to trigger stress-induced activation of HSPs (Baird *et al*. 2014). Moreover, transgenic HSF-1 activation promotes survival in a neurodegenerative mouse model without inducing increased expression of HSPs in brain tissue (Fujimoto *et al*. 2005). These findings suggest that life span extension associated with increased HSF-1 activity in animals is not solely due to upregulation of canonical HSR genes, but it presumably also involves transcriptional regulation of other unidentified HSF-1 targets. In addition to its role in HSR, HSF-1 has major functions in other biological processes such as development, reproduction, metabolism and cancer (Li *et al*. 2017). Hence, increasing the gene dosage of *hsf-1* might ectopically affect the expression of a large number of HSF-1 target genes that are not directly involved in regulation of longevity in animals.

In normal physiological conditions, the transactivation potential of HSF-1 is limited by several regulatory mechanisms that dictate the context-dependent activation status of the HSF-1 protein (Anckar and Sistonen 2011; Gomez-Pastor *et al*. 2018). One such negative regulator of HSF-1 is the evolutionarily conserved heat shock factor binding protein 1 (HSB-1) (Morimoto 1998). Direct interaction between the human homologs of HSB-1 and HSF-1 in a yeast two-hybrid screen suggested that binding of HSB-1 to the trimerization domain of HSF-1 can inhibit its transactivation potential *in vivo* (Satyal *et al*. 1998). In *C*. *elegans*, HSB-1 physically binds to HSF-1 to form an inhibitory multiprotein complex (Satyal *et al*. 1998; Chiang *et al*. 2012). Interestingly, the formation of this HSF-1-inhibitory complex is not affected by heat stress, but instead is promoted by insulin/IGF-1-like signaling (Chiang *et al*. 2012), an evolutionarily conserved longevity regulating pathway (Riera *et al*. 2016). Genetic ablation of *hsb-1* results in dissociation of HSF-1 from this inhibitory complex and induces a robust increase in life span of animals that is dependent on HSF-1 activity (Chiang *et al*. 2012). However, it remains elusive how the absence of HSB-1 alters the transactivation potential of HSF-1, and thus promotes organismal longevity via potentially modifying the expression of certain HSF-1 target genes. We hypothesized that inhibition of HSB-1 influences a small subset of the HSF-1-regulated transcriptome that has a specific role in regulation of longevity in *C*. *elegans* (Figure 1). In contrast, increasing the gene dosage of *hsf-1* potentially induces transcriptional changes for a vast majority of HSF-1 target genes involved in varied biological processes. A previous study established that a sizeable proportion of the entire *C*. *elegans* transcriptome comprises HSR-independent downstream targets of HSF-1 (Brunquell *et al*. 2016). Surprisingly, expression levels of most of the HSF-1-regulated genes identified in that study were not found to be significantly altered during normal aging in *C*. *elegans* (Budovskaya *et al*. 2008; Brunquell *et al*. 2016). This further indicates that studying the transcriptional changes associated with inhibition of HSB-1 can help in narrowing down the HSF-1 target genes that are essential for mediating the HSR-independent longevity-promoting effects of HSF-1. Though homologs of *C*. *elegans hsb-1* exist in genomes of all multicellular eukaryotes (Morimoto 1998), the function of this gene in normal physiological conditions remains less explored in complex organisms (Dirks *et al*. 2010; Eroglu *et al*. 2014).

**Figure 1.**
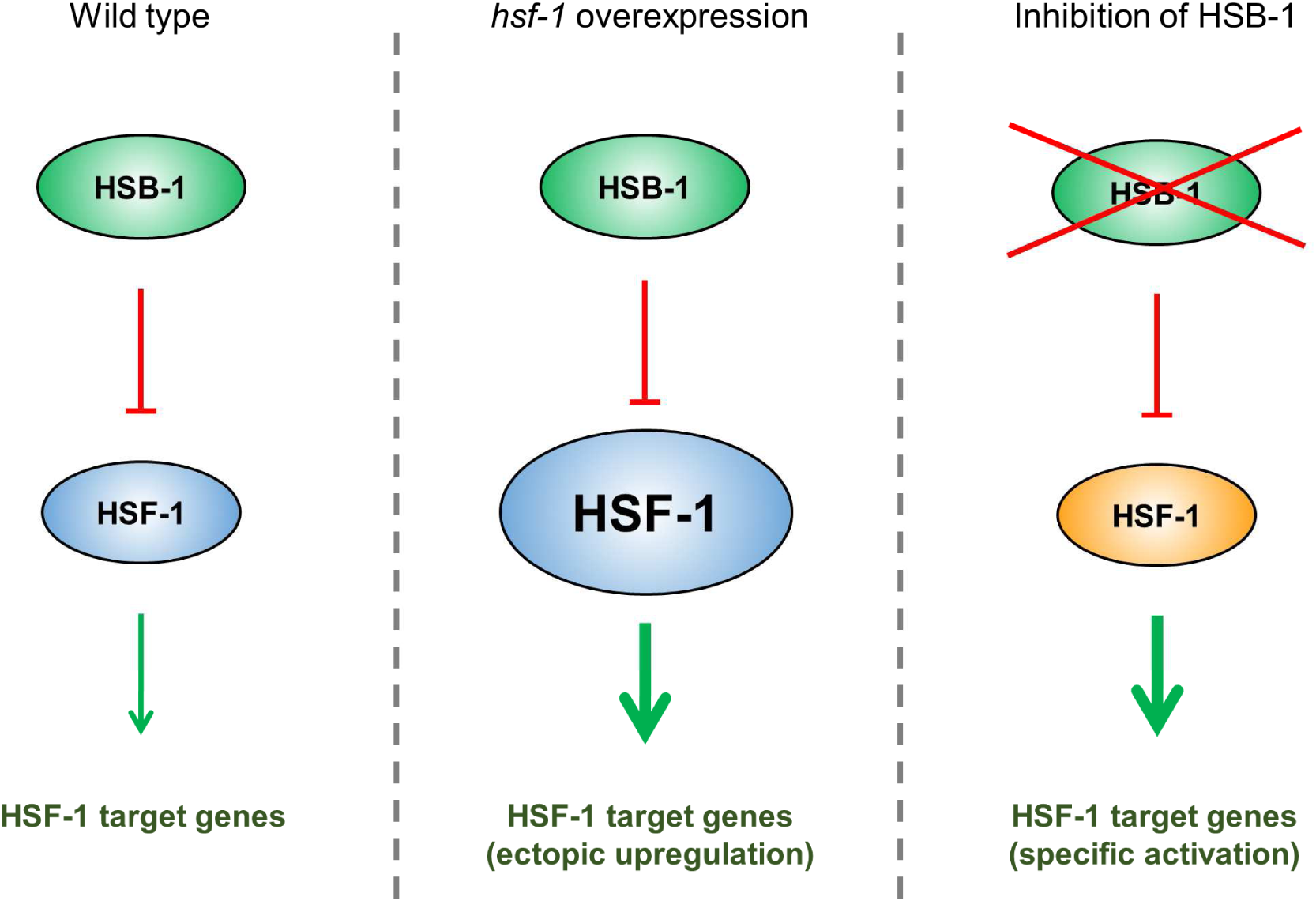
Proposed hypothesis of specific regulation of HSF-1 target genes by HSB-1. In wild-type animals, HSB-1 binds to HSF-1 and limits its transactivation potential (*left*). *hsf-1* overexpression alters transcript levels of a majority of HSF-1 target genes. In addition to their effect on longevity, these genes also modulate other pathways such as stress response, development and reproduction (*middle*). We hypothesize that in the absence of HSB-1, HSF-1 acquires an altered transactivation potential (shown in orange), which leads to a selective change in the HSF-1-regulated transcriptome that extends organismal life span (*right*).

Here we show that in the absence of HSB-1 regulation, heat shock factor acquires an altered transactivation profile, which results in extensive changes to the *C*. *elegans* transcriptome. Genetic ablation of *hsb-1* induces increased *in vitro* DNA binding activity of the HSF-1 transcription factor similar to that observed in an HSF-1 overexpression strain, even though HSF-1 protein level is not elevated in the *hsb-1* mutant. Roughly 45% of the genes that are differentially expressed in the *hsb-1(-)* strain also constitute a subset of the altered transcriptome in *hsf-1* overexpressing animals, but the global transcriptional changes associated with HSB-1 inhibition are distinct from that induced via overexpression of HSF-1. We further show that the subset of genes that are differentially expressed in both *hsb-1(-)* and *hsf-1* overexpressing animals have a strongly correlated expression pattern in the two strains. Overall, our findings collectively suggest that HSB-1 controls the expression of a small subset of the HSF-1-regulated transcriptome, which might be essential for mediating the longevity promoting effects of HSF-1 in normal physiological conditions.

## MATERIALS AND METHODS

### *C*. *elegans* Strains

All *C*. *elegans* strains used in this study were maintained at 20ºC on nematode growth medium (NGM) plates seeded with *E*. *coli* OP50 strain using the standard method. Animals were maintained for at least three generations without starvation before they were used for experiments. The following strains were used in this study: N2: Wild-type, EQ150: *hsb-1(cg116) IV* – CH116 outcrossed 4x to wild-type N2, and EQ140: *iqIs37*[pAH76(*hsf-1p∷myc-hsf-1*) + pRF4(*rol-6p∷rol-6(su1006)*)]. Wild-type (N2) and CH116 strains were obtained from *Caenorhabditis* Genetics Center (Saint Paul, MN). All experiments in this study were performed using day 1 adult animals that were obtained after age-synchronization of unhatched eggs.

### RNA Isolation and Quantitative RT-PCR

∼2,000 age-synchronized young adults grown on NGM plates were collected in M9 buffer containing 0.01% Triton X-100 and were immediately washed three times with M9 buffer. TRIzol reagent (Thermo Fisher) was used to extract total RNA using the manufacturer’s protocol and TURBO DNA-free kit (Themo Fisher) was used to perform DNase-treatment. cDNA was synthesized from 5 µg RNA with 0.5 µg of oligo(dT)_12-18_ primer (Thermo Fisher) using SuperScript III reverse transcriptase (Thermo Fisher). Quantitative PCR was performed with Power SYBR Green PCR master mix (Thermo Fisher) and PCR primers listed in Table S3 using a CFX96 Real-Time PCR detection system (Bio-Rad). Primers were designed using the Primer3Plus online tool. Comparative Ct method was used to measure relative transcript levels while using the housekeeping gene *cdc-42* as an internal control for normalization (Hoogewijs *et al*. 2008). Normalization of transcript levels using another housekeeping gene *pmp-3* led to similar results (Hoogewijs *et al*. 2008; Zhang *et al*. 2012).

### Immunoblot Analysis

∼10,000 age-synchronized young adults grown on HG plates were collected in M9 buffer containing 0.01% Triton X-100 and were immediately washed three times with M9 buffer. Pellets were suspended in three times the volume of 6x Sample Buffer (375 mM Tris-Cl at pH 6.8, 12% SDS, 60% glycerol, 600 mM DTT, 0.06% Bromophenol Blue) and were snap-frozen in liquid nitrogen for 30 secs. Samples were subsequently boiled for 10 min, vortexed for 15 sec, centrifuged at 16,000 × g for 30 min and subjected to SDS-PAGE on a SE 250 mini-vertical unit (GE Healthcare). Proteins were transferred to Immobilon-P PVDF membrane (Millipore Sigma) at 400 mA for 40 min on a Trans-blot SD semi-dry transfer cell (Bio-Rad) and the PVDF membrane was blocked in 5% non-fat dry milk-based blocking-grade blocker (Bio-Rad) for 1 hr at room temperature. Primary antibody incubation was performed for 12–16 hrs at 4ºC using the following antibodies: HSF-1 – Abnova custom-made (1:1,000) lot no. F4271-3E11 and β-actin – Abcam ab8227 (1:10,000). The PVDF membrane was washed four times with TBS buffer containing 0.1% Tween 20 (TBS-T) for 5 min per wash, incubated with HRP-conjugated secondary antibody for 1 hr at room temperature and washed again four times with TBS-T buffer for 5 min per wash. Blots were developed using Immobilon Western Chemiluminescent HRP Substrate (Millipore Sigma) and visualized by autoradiography.

### Preparation of Nuclear Extracts

∼4,000 age-synchronized young adults grown on NGM plates were collected in M9 buffer containing 0.01% Triton X-100 and were immediately washed three times with M9 buffer. Samples were frozen in liquid nitrogen and thawed on ice for two freeze-thaw cycles. Pellets were resuspended in 10 times the volume of NPB buffer (15 mM HEPES-Na at pH 7.6, 10 mM KCl, 1.5 mM MgCl_2_, 0.1 mM EDTA, 0.5 mM EGTA, 44 mM sucrose, 1 mM dithiothreitol, protease inhibitors, phosphatase inhibitors) containing 0.25% NP-40 and 0.1% Triton X-100. Animals were homogenized by 20 strokes of pestle B of the Kontes Dounce tissue grinder. The homogenized samples were centrifuged at 4,000 × g for 5 min and the nuclear pellet was washed two times with NPB buffer containing 0.25% NP-40 and 0.1% Triton X-100. Nuclei were extracted in four times the volume of HEG buffer (20 mM HEPES-Na at pH 7.9, 0.5 mM EDTA, 10% glycerol, 420 mM NaCl, 1.5 mM MgCl_2_, and protease inhibitors) at 4°C for 1 hr. The nuclear fraction was collected by centrifugation at 14,000 × g for 15 min. Protein concentrations were determined by Bradford assay.

### Electrophoretic Mobility Shift Assay

Binding of HSF-1 to its genomic target sequence was measured by Electrophoretic Mobility Shift Assay (EMSA) using a previously described protocol (Chiang *et al*. 2012). Briefly, 1 μg of nuclear extract was incubated for 20 min at room temperature in EMSA binding buffer (10 mM Tris-Cl at pH 7.5, 50 mM KCl, 1 mM DTT) supplemented with 50 ng/μL Poly (dI.dC) and 1 nM biotin-TEG-labeled oligonucleotide containing an HSE sequence. The biotin-TEG-labeled oligonucleotide was synthesized by annealing two complementary sequences (listed in Table S3) corresponding to an HSE from the promoter region of *hsp-16*.*1* gene. The DNA-bound nuclear extracts were resolved using native 3.5% polyacrylamide gel electrophoresis and were transferred to Immobilon-Ny+ charged nylon membrane (Millipore Sigma) at 400 mA for 1 hr on a Trans-blot SD semi-dry transfer cell (Bio-Rad). The HSF-1-HSE DNA complexes were visualized by autoradiography using LightShift Chemiluminescent EMSA kit (Thermo Fisher).

### RNA-Seq Library Preparation

TRIzol reagent (Thermo Fisher) was used to extract total RNA from ∼1,000 age-synchronized young adults of N2, EQ150 and EQ140 strains (three biological replicates per genotype) using the manufacturer’s protocol and DNase-treatment was performed using TURBO DNA-free kit (Themo Fisher). mRNA was isolated from the DNase-treated total RNA using oligo(dT) and cDNA was synthesized by reverse transcription using TruSeq RNA Library Prep Kit v2 (Illumina). The samples were subsequently end-repaired, poly(A)-tailed, ligated to barcoded oligo adapters (Illumina) and PCR-amplified using Apollo 324 library preparation system (Wafergen Bio-systems) to prepare non-stranded poly(A)-based cDNA libraries with 120 bp insert size. All samples were multiplexed for 50 bp single-end sequencing on a HiSeq 2500 platform (Illumina).

### RNA-Seq Data Analysis

Quality control and adaptor trimming of raw sequencing reads were performed using Trim Galore version 0.4.5 (available at https://github.com/FelixKrueger/TrimGalore) (Martin 2011). Trimmed RNA-Seq reads were mapped to the *C*. *elegans* reference transcriptome (WS260 release) using STAR version 2.6.0c (Dobin *et al*. 2013). Subsequently, transcript abundance of each gene was measured using HTSeq version 0.9.1 (Anders *et al*. 2015). To identify genes that were differentially expressed relative to wild type, transcript counts were analyzed by DESeq2 version 1.20.0 (Love *et al*. 2014). Genes with read counts less than 10 reads were excluded for the DESeq2 analysis. In total, 15,787 genes were assayed for differential expression analysis between strains, using the following two criteria: false discovery rate (FDR)-adjusted p value < 0.05 and fold-change ≥ 1.5. For principal component analysis (PCA) and pairwise distance analysis, counts with variance stabilizing transformation (VST) were used. Venn diagrams for differentially expressed genes were generated using Venn Diagram Plotter (Pacific Northwest National Laboratory; available at https://omics.pnl.gov/software/venn-diagram-plotter). For each genotype, the genes differentially expressed relative to wild-type were annotated with Gene Ontology (GO) terms from NCBI and the Database for Annotation, Visualization and Integrated Discovery (DAVID) was used to identify significantly enriched functional categories from the overrepresented GO terms (Huang *et al*. 2009). To determine the proportion of known longevity genes among the differentially expressed genes identified in this study, phenotype data for genes were accessed from https://www.wormbase.org. Specifically, a gene was classified as a longevity gene if it is annotated with one or more of these phenotype terms in Wormbase: ‘shortened life span’, ‘extended life span’ and ‘life span variant’.

### Statistical Analysis

GraphPad Prism 8 was used to perform statistical analyses and graphing. Hypergeometric tests were performed in R. Details of statistical tests used, number of biological replicates and *p* values for each experiment are included in the figure legends.

### Data and Reagent Availability

Raw RNA-Seq reads have been deposited at the NCBI Gene Expression Omnibus (GEO) under the accession number GSE119993. Supplemental materials uploaded to the GSA figshare portal include: list of differentially expressed genes in *hsb-1(-)* mutant and *hsf-1* overexpression strains (Tables S1 and S2, respectively); and DNA sequences of oligonucleotide primers used in EMSA and quantitative PCR experiments (Table S3). All *C*. *elegans* strains used in this study are available upon request.

## RESULTS

### HSF-1 acquires increased transactivation potential in the absence of HSB-1 regulation that is distinct from heat stress-induced HSF-1 activation

Previous studies have shown that HSF-1-dependent life span extension can be achieved in *C*. *elegans* via increasing the gene dosage of *hsf-1* or genetic ablation of *hsb-1* (Hsu *et al*. 2003; Morley and Morimoto 2004; Chiang *et al*. 2012). To elucidate whether inhibition of HSB-1 promotes longevity in animals via increasing *hsf-1* expression, we compared the transcript levels of *hsf-1* in wild-type (N2), *hsb-1(-)* and a previously described long-lived *hsf-1* overexpression strain (Chiang *et al*. 2012). Surprisingly, unlike in the *hsf-1* overexpression strain, the transcript levels of *hsf-1* were not significantly different in *hsb-1(-)* animals compared to wild-type (Figure 2A). Furthermore, while the protein level of HSF-1 was elevated by roughly 60% in the *hsf-1* overexpression strain, *hsb-1(-)* animals did not have increased HSF-1 protein level compared to wild-type (Figure 2B). This indicated that though HSB-1 inhibition has been shown to extend *C*. *elegans* life span in an *hsf-1*-dependent manner (Chiang *et al*. 2012), it does not involve an increase in expression of *hsf-1* in the long-lived *hsb-1(-)* strain. Incidentally, HSF-1 activation in the presence of heat stress also does not involve transcriptional upregulation of *hsf-1* (Chiang *et al*. 2012), but instead necessitates several post-translational modifications to the HSF-1 protein (Anckar and Sistonen 2011; Chiang *et al*. 2012; Gomez-Pastor *et al*. 2018). Since HSB-1 is known to bind to HSF-1 and form an inhibitory multiprotein complex (Satyal *et al*. 1998; Chiang *et al*. 2012), we speculated that the absence of HSB-1 might result in release of HSF-1 from this inhibitory complex and thus increase its transcriptional activation potential. Previous findings have shown that genetic ablation of *hsb-1* prevents the formation of this HSF-1-inhibitory complex (Chiang *et al*. 2012). We used EMSA to investigate if the DNA binding activity of HSF-1 to its genomic target sequence is altered in the absence of HSB-1 regulation. In nuclear extracts from wild-type animals subjected to heat stress, the *in vitro* binding activity of HSF-1 to its target HSE sequence increased by greater than 2-fold compared to that from non-stressed animals (Figure 2C), as shown previously (Chiang *et al*. 2012). Interestingly, in non-stressed basal conditions, HSF-1 from nuclear extracts of both *hsb-1(-)* and *hsf-1* overexpressing animals showed ∼80% increase in *in vitro* HSE binding activity compared to that in wild-type animals (Figure 2C). This suggests that even in non-stressed conditions, HSB-1 inhibition and HSF-1 overexpression can potentially promote increased HSF-1 binding to its genomic target sequences, similar to that observed during heat stress.

**Figure 2.**
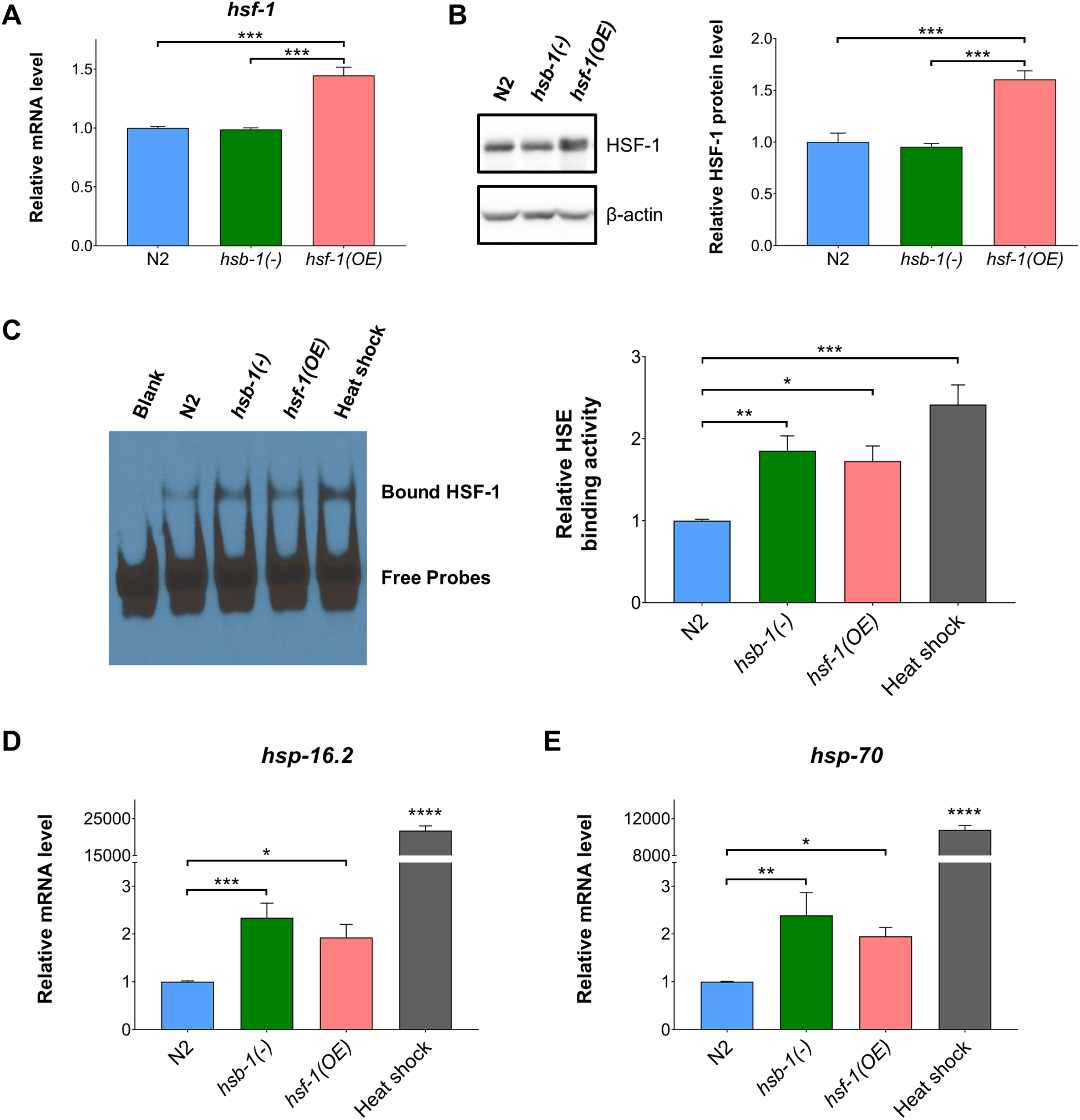
HSB-1 inhibition results in higher HSF-1 activity without involving an increase in HSF-1 protein level. **(A)** Relative transcript levels of *hsf-1* in wild-type (N2), *hsb-1(-)* and *hsf-1* overexpression strains grown at 20°C. All values are reported in comparison to mean wild-type expression level in control conditions. Data represent mean ± SEM for ≥ 3 biological replicates. *** *p* < 0.001 in Tukey’s multiple comparisons test performed after one-way ANOVA. **(B)** Representative immunoblots for HSF-1 and β-actin proteins in N2, *hsb-1(-)* and *hsf-1* overexpression strains grown at 20°C (*left*). Densitometric quantification of HSF-1: β-actin ratio relative to wild-type levels (*right*). Data represent mean ± SEM for ≥ 3 biological replicates. *** *p* < 0.001 in Tukey’s multiple comparisons test performed after one-way ANOVA. **(C)** Representative EMSA analysis for HSE binding activity of nuclear extracts from N2, *hsb-1(-)* and *hsf-1* overexpression strains grown in non-stressed conditions at 20°C and N2 animals subjected to heat stress at 37°C for 90 min (*left*). Densitometric quantification of HSE binding activity relative to wild-type levels (*right*). Data represent mean ± SEM for ≥ 4 biological replicates. * *p* < 0.05, ** *p* < 0.01 and *** *p* < 0.001 in Tukey’s multiple comparisons test performed after one-way ANOVA. **(D, E)** Relative transcript levels of *hsp-16*.*2* and *hsp-70* in N2, *hsb-1(-)* and *hsf-1* overexpression strains grown in non-stressed conditions at 20°C and N2 animals subjected to heat stress at 37°C for 90 min. Data represent mean ± SEM for ≥ 3 biological replicates. * *p* < 0.05, ** *p* < 0.01 and *** *p* < 0.001 in Tukey’s multiple comparisons test performed after one-way ANOVA. **** indicates *p* < 0.0001 compared to all other conditions in Tukey’s multiple comparisons test.

Next, we tested if HSB-1 inhibition and HSF-1 overexpression trigger HSF-1-mediated transcriptional changes in animals, as observed during heat stress. We measured transcript levels of two canonical HSF-1 target genes, *hsp-16*.*2* and *hsp-70*, that are upregulated during heat stress in *C*. *elegans* (GuhaThakurta *et al*. 2002; Chiang *et al*. 2012). The transcript levels of both *hsp-16*.*2* and *hsp-70* genes increased by several thousand-fold after animals were subjected to 90 min of heat shock at 37°C (Figures 2D and 2E). Intriguingly, both HSB-1 inhibition and HSF-1 overexpression resulted in a significant ∼2-fold increase in transcript levels of both these *hsp* genes relative to wild-type levels in non-stressed conditions (Figures 2D and 2E). These findings collectively indicate that though heat stress, HSB-1 inhibition and HSF-1 overexpression all induce an increase in the *in vitro* DNA binding activity of HSF-1 (Figure 2C), the latter two conditions have a widely distinct effect on the transcription profile of canonical HSR genes compared to that observed during heat stress (Figures 2D and 2E). Hence, the nature of HSF-1 activation resulting due to overexpression of HSF-1 or inhibition of its negative regulator HSB-1 are possibly distinct from its broadly studied heat stress-induced activation (Li *et al*. 2017; Gomez-Pastor *et al*. 2018).

### HSF-1 overexpression, but not HSB-1 inhibition, induces large-scale transcriptional upregulation in *C*. *elegans*

HSF-1 target genes that mediate its longevity-promoting effects in animals still remain largely unidentified. Since both HSB-1 inhibition and HSF-1 overexpression have been previously reported to induce life span extension via increasing the activity of the HSF-1 transcription factor (Hsu *et al*. 2003; Morley and Morimoto 2004; Chiang *et al*. 2012), we speculated that these two manipulations might induce similar changes to the HSF-1-regulated transcriptome in *C*. *elegans*. For this comparison, we performed RNA-Seq to evaluate the gene expression profiles in N2, *hsb-1(-)* and *hsf-1* overexpression strains (Figure 3A). At the whole transcriptome level, within-group variation among the three biological replicates for each genotype was lesser than between-group variation across genotypes (Figures 3B and 3C). This indicates that both HSB-1 inhibition and HSF-1 overexpression induce highly reproducible gene expression changes in animals. Strikingly, the transcriptome-wide expression pattern in the *hsb-1(-)* strain was more similar to that of wild-type animals, rather than to the expression pattern in the *hsf-1* overexpression strain (Figure 3C).

**Figure 3.**
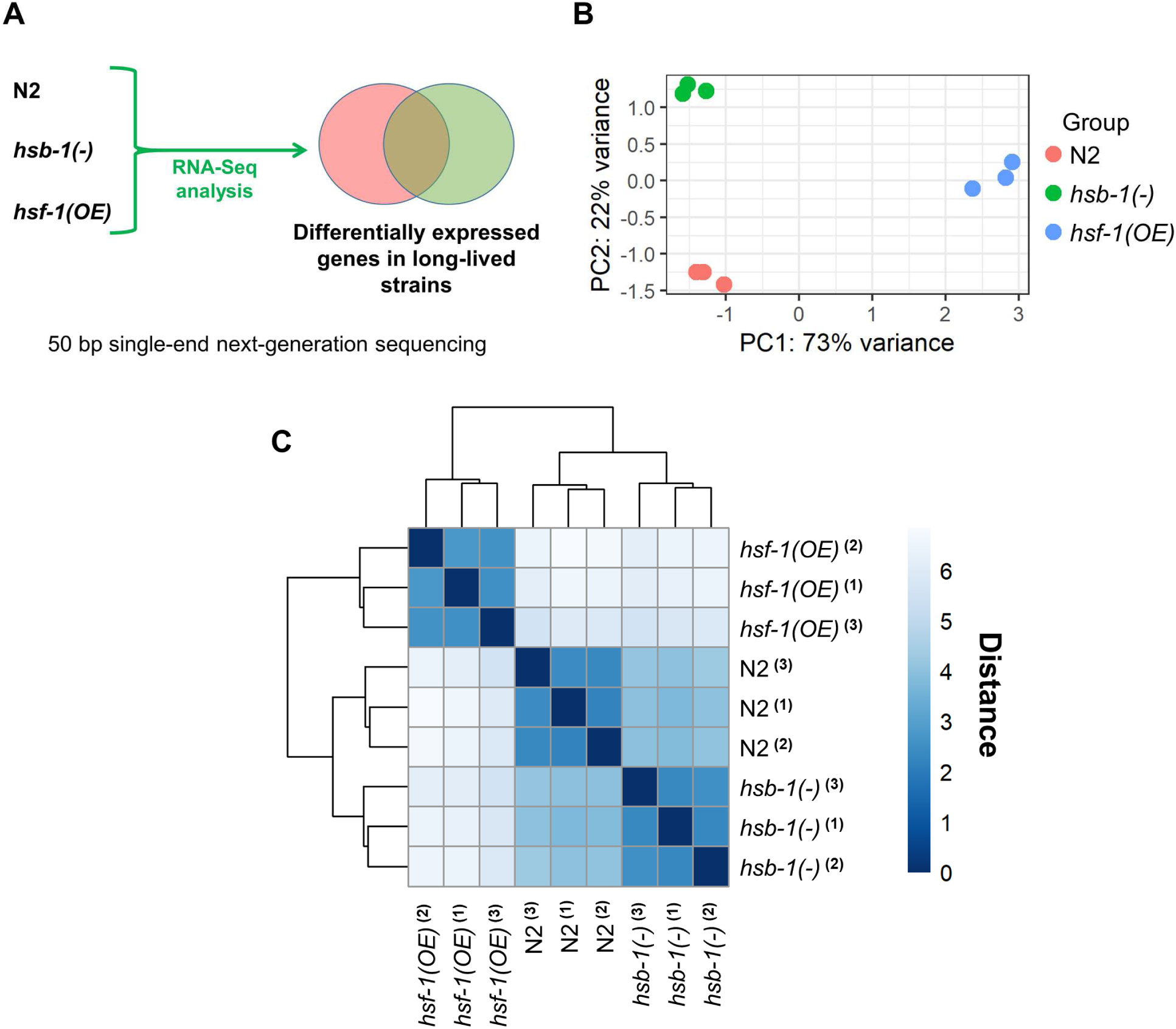
RNA-Seq analysis for comparison of the transcriptomes of *hsb-1(-)* and *hsf-1* overexpressing animals. **(A)** Schematic of RNA-Seq study design for identifying differentially expressed genes in *hsb-1(-)* and *hsf-1* overexpression strains relative to wild-type (N2). **(B)** Principal component analysis (PCA) plot showing variability between all biological replicates at the whole-transcriptome level. **(C)** Pairwise distance heatmap showing similarities between biological replicates and genotypes based on transcriptome-wide expression pattern.

Next, we generated MA plots and volcano plots to visualize genes for which the fold-change in expression relative to wild-type was statistically significant with an FDR-adjusted *p* value of less than 0.05 (Figures 4A–D). 954 transcripts were found to have a statistically significant difference in expression in *hsb-1(-)* animals (Figures 4A and 4C), while 5,144 transcripts had significantly different expression in the *hsf-1* overexpression strain (Figures 4B and 4D). The MA and volcano plots also indicated that HSF-1 overexpression induces large-scale transcriptional changes for a large number of *C*. *elegans* genes (Figures 4B and 4D). In addition to the criterion of statistical significance, we set a threshold of at least 1.5-fold change in transcript levels relative to wild-type to identify gene expression changes that are potentially of biological relevance. Genes that were ≥1.5-fold (±0.585 on log_2_ scale) upregulated or downregulated with an FDR-adjusted *p* value of less than 0.05 were identified as being differentially expressed. With these two criteria, we found 1,662 genes to be differentially expressed in the *hsf-1* overexpression strain (Figure 5A and Table S2). This was in agreement with previous studies showing thousands of genes as being potential transcriptional targets of HSF-1, even in non-stressed physiological conditions (Mendillo *et al*. 2012; Brunquell *et al*. 2016). More than 95% of these 1,662 genes had upregulated expression in *hsf-1* overexpressing animals, while the remaining less than 5% were downregulated (Figures 5B and 5C). In contrast, we found only 477 genes to be differentially expressed in the *hsb-1(-)* strain, though interestingly, roughly 45% of these overlapped with the differentially expressed gene list in the *hsf-1* overexpression strain (Figure 5A, and Tables S1 and S2). In *hsb-1(-)* animals, the pattern of differential expression was slightly different to that in *hsf-1* overexpressing animals, with ∼78% of genes being upregulated and remaining ∼22% being downregulated (Figures 5B and 5C). In addition, more than 46% of genes that were upregulated in the *hsb-1(-)* strain also showed significant upregulation in the *hsf-1* overexpression strain, but this overlap was less than 18% for downregulated genes (Figures 5B and 5C). These findings indicate a considerable overlap between the upregulated transcriptomes associated with HSB-1 inhibition and HSF-1 overexpression in *C*. *elegans*.

**Figure 4.**
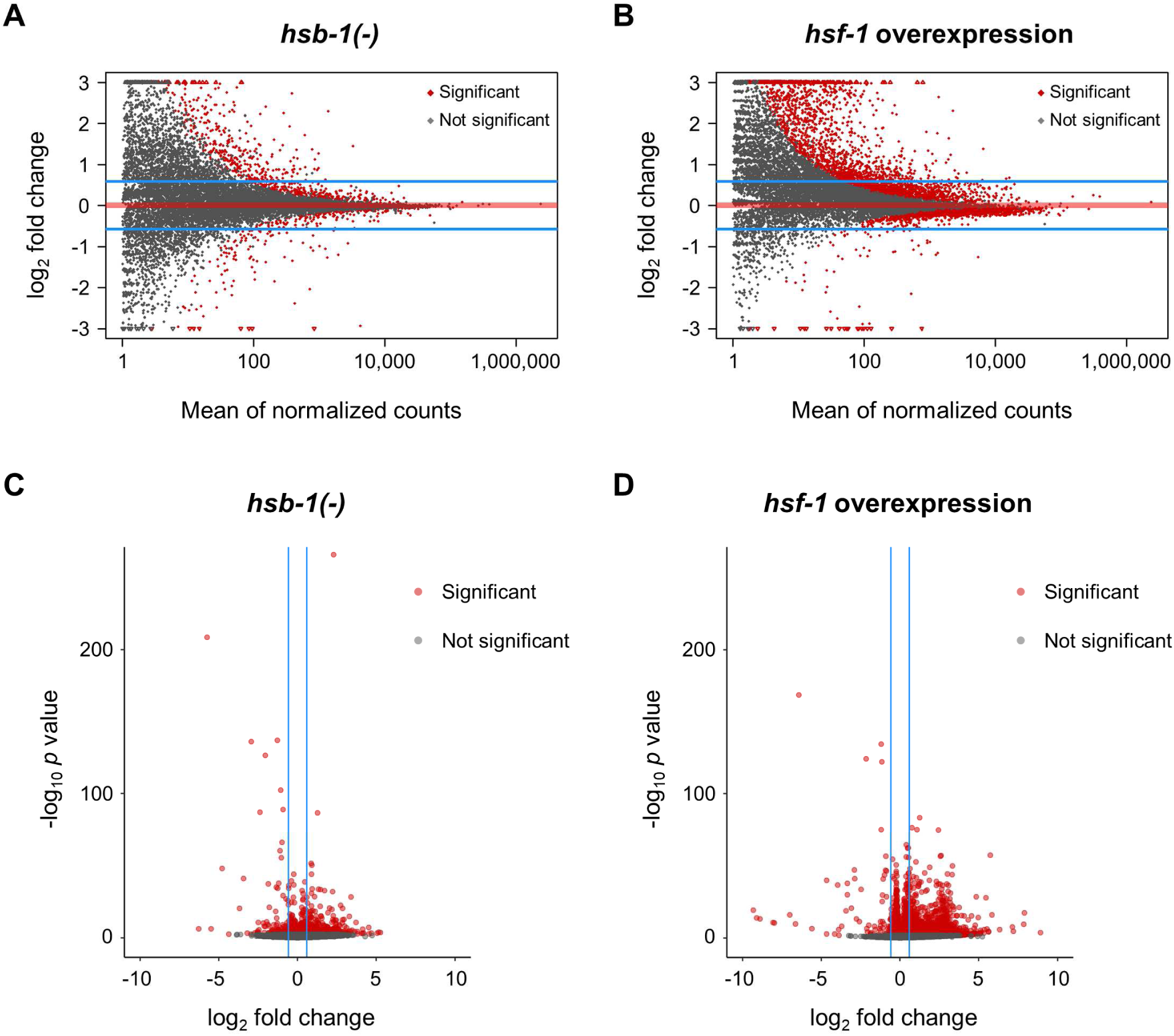
Global transcriptional profile indicates large-scale gene expression changes in *hsf-1* overexpressing animals. **(A, B)** MA plots for all genes with significant (FDR-adjusted *p* < 0.05) or no significant change in expression in *hsb-1(-)* and *hsf-1* overexpression strains relative to wild-type (N2) arranged in order of mean expression level. Read counts for genes were normalized to the size factors determined from DESeq2. Horizontal blue lines encompass genes with less than 1.5-fold change in expression (±0.585 on log_2_ scale) relative to N2. Horizontal red line represents no change in expression. **(C, D)** Volcano plots for all genes with statistically significant (FDR-adjusted *p* < 0.05) or no significant change in expression in *hsb-1(-)* and *hsf-1* overexpression strains relative to wild-type levels. Vertical blue lines encompass genes with less than 1.5-fold change in expression (±0.585 on log_2_ scale) relative to N2.

**Figure 5.**
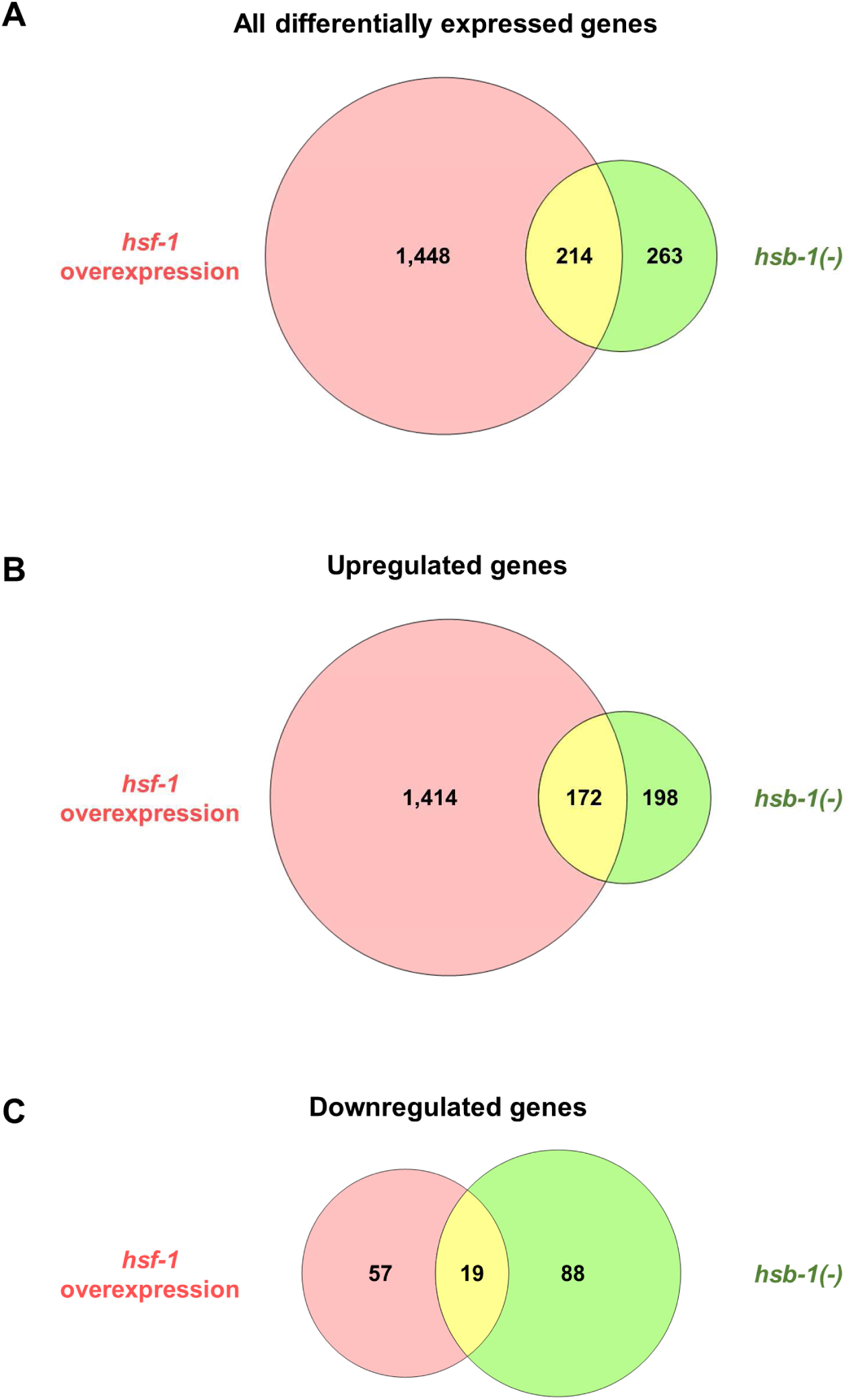
Venn diagrams for differentially expressed genes in *hsb-1(-)* and *hsf-1* overexpression strains. **(A–C)** Venn diagrams showing number of differentially expressed genes (fold-change ≥ 1.5) only in *hsb-1(-)* animals, only in *hsf-1* overexpressing animals or in both the strains relative to wild-type levels. **(B)** shows number of upregulated genes, **(C)** shows number of downregulated genes, while **(A)** shows number of genes in either of the two categories.

### HSB-1 inhibition and HSF-1 overexpression induce differential expression of genes that represent overlapping biological processes

To identify the biological processes that are potentially affected by the altered transcriptome in *hsb-1(-)* and *hsf-1* overexpressing animals, we used pathway enrichment analysis using GO databases to determine functional categories associated with the genes that were differentially expressed in these two strains. Inhibition of HSB-1 and overexpression of HSF-1 affected expression of genes representing overlapping biological functions (Figures 6A and 6B). Some GO terms, such as pseudopodium (corresponding to genes involved in sperm motility), major sperm protein, PapD-like superfamily, acetylation and cell projection were represented by differentially expressed genes in both *hsb-1(-)* and *hsf-1* overexpression strains (Figures 6A and 6B). PapD-like superfamily in *C*. *elegans* constitutes major sperm proteins, which contain an immunoglobin-like domain structurally similar to PapD-like chaperones. Moreover, in the list of differentially expressed genes identified in our RNA-Seq comparison, the GO terms for acetylation and cell projection were primarily represented by multiple major sperm protein (*msp*) family genes that were found to be significantly upregulated in both *hsb-1(-)* and *hsf-1* overexpression strains (Tables S1 and S2). Previous studies performed in mice have indicated an essential role of *Hsf1* and *Hsf2* genes in spermatogenesis (Nakai *et al*. 2000; Wang *et al*. 2004). Mice expressing a constitutively active form of the HSF1 protein show altered expression of several genes involved in sperm production (Nakai *et al*. 2000). Since several differentially expressed genes in both *hsb-1(-)* and *hsf-1* overexpression strains in *C*. *elegans* represented multiple GO terms corresponding to sperm function (Figures 6A and 6B), we speculate that this is due to an evolutionarily conserved role of the HSF-1 transcription factor in spermatogenesis (Abane and Mezger 2010). Moreover, the GO term for cytoskeleton was enriched in the differentially expressed gene list for both *hsb-1(-)* and *hsf-1* overexpression strains (Figures 6A and 6B), which corroborates the recently discovered role of cytoskeletal components in HSF-1-induced longevity mechanisms (Baird *et al*. 2014). In addition, both the long-lived strains showed elevated expression of several genes involved in the innate immune response (Figures 6A and 6B), a pathway that has been previously implicated in longevity regulation in *C*. *elegans* (Yunger *et al*. 2017). The gene encoding for the Toll/interleukin-1 receptor adaptor protein TIR-1, an essential component of the innate immune response in nematodes (Liberati *et al*. 2004), was found to be significantly upregulated in both *hsb-1(-)* and *hsf-1* overexpression strains (Tables S1 and S2). Interestingly, a recent study has shown a major role of *tir-1* in mediating the life span extension associated with activation of p38 MAPK pathway in *C*. *elegans* (Verma *et al*. 2018).

**Figure 6.**
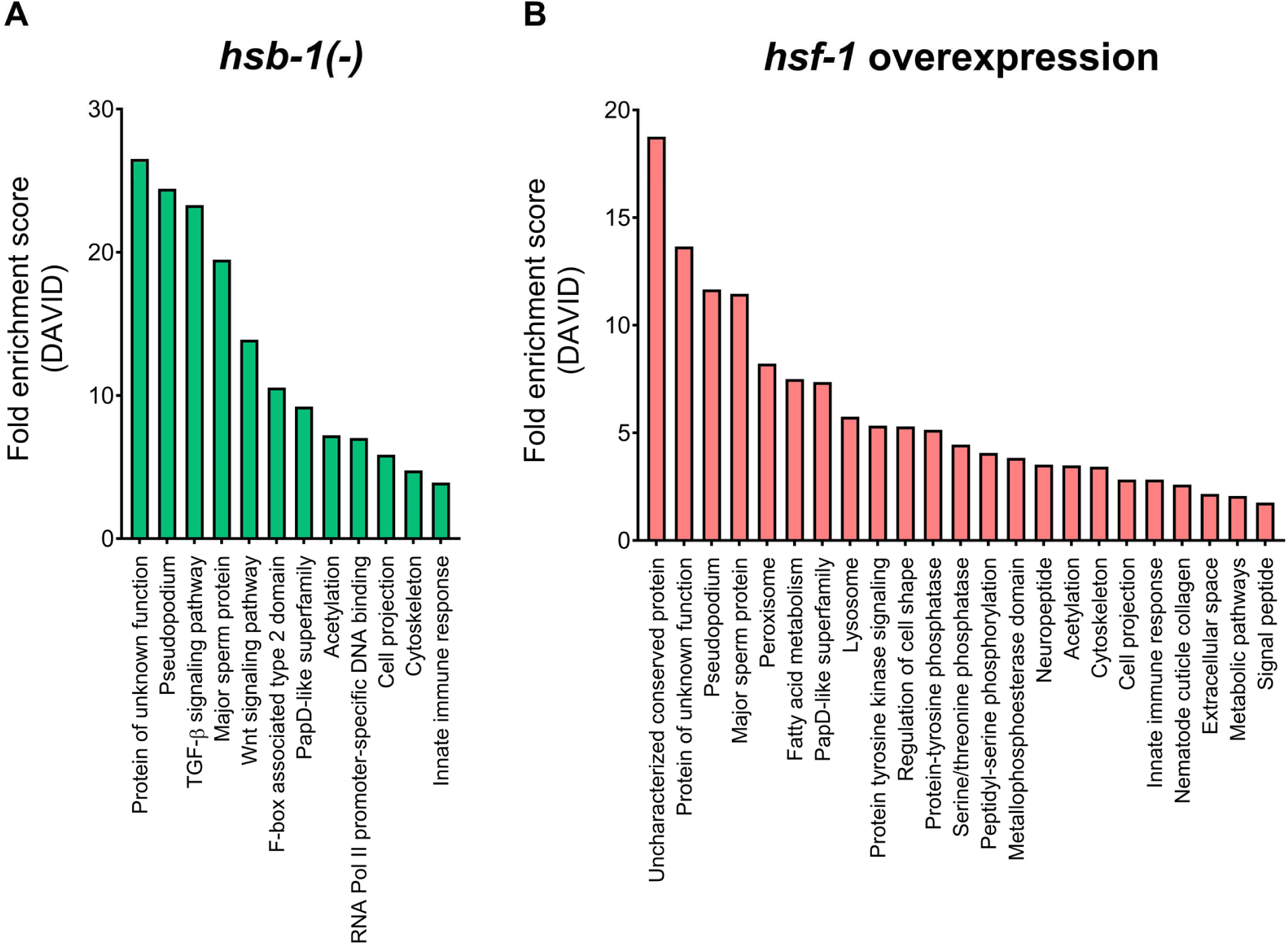
Gene ontology (GO) analysis of differentially expressed genes in *hsb-1(-)* and *hsf-1* overexpression strains. **(A, B)** Functional enrichment analysis (DAVID) of differentially expressed genes in *hsb-1(-)* and *hsf-1* overexpression strains relative to wild-type. GO terms representing unique functional categories with FDR-adjusted *p* < 0.05 are shown.

In contrast, certain functional categories were found to be unique to each genotype. Genes differentially expressed due to HSF-1 overexpression were associated with peroxisomal function (Figure 6B). The peroxisomal catalase gene *ctl-2* was significantly upregulated in *hsf-1* overexpressing animals, but not in the *hsb-1(-)* mutant (Tables S1 and S2). Previous studies have shown that *ctl-2* activity is required for the longevity induced via reduced insulin/IGF-1-like signaling in *C*. *elegans* (Murphy *et al*. 2003). On the other hand, genes differentially expressed due to HSB-1 inhibition were associated with TGF-β and Wnt signaling pathways (Figure 6A), processes that have been shown to have a direct role in longevity regulation in *C*. *elegans* (Shaw *et al*. 2007; Lezzerini and Budovskaya 2014). GO terms for these two pathways were enriched due to *hsb-1(-)*-specific upregulation of multiple members of the Skp1-related (*skr*) gene family (Table S1). The *skr* genes encode components of the proteasomal E3 ubiquitin ligase complex and have a role in mediating life span extension in insulin/IGF-1-like signaling mutants (Ghazi *et al*. 2007). Overall, our pathway enrichment analyses suggested that HSB-1 inhibition and HSF-1 overexpression affect mostly overlapping but also certain unique biological processes in *C*. *elegans*.

### DAF-16 targets that are upregulated via both HSB-1 inhibition and HSF-1 overexpression include numerous known longevity genes

Next, we compared the list of differentially expressed genes identified in this study with a previously reported dataset for transcriptional changes associated with insulin/IGF-1-like signaling in *C*. *elegans* (Tepper *et al*. 2013). Reduced activity of the insulin/IGF-1-like receptor DAF-2 is one of the first reported and most widely-studied longevity pathways in *C*. *elegans* (Kenyon *et al*. 1993). Life span extension in the loss-of-function *daf-2* mutant is mediated via increased activity of the FOXO transcription factor DAF-16 (Kenyon *et al*. 1993; Ogg *et al*. 1997; Lin *et al*. 1997). Interestingly, HSF-1 activity is also upregulated in the *daf-2* mutant (Chiang *et al*. 2012), and the longevity phenotype of this strain can be completely suppressed via knockdown of *hsf-1* (Hsu *et al*. 2003). Numerous studies have indicated cooperation between DAF-16 and HSF-1 to mediate life span extension in *C*. *elegans* (Hsu *et al*. 2003; Cohen *et al*. 2006; Douglas *et al*. 2015), though the nature of interaction between these two pro-longevity transcription factors remains largely unknown. Here, we compared the genes found to be differentially expressed via HSB-1 inhibition or HSF-1 overexpression in *C*. *elegans* with previously reported DAF-16 target genes that have altered expression in *daf-2* mutant animals (Tepper *et al*. 2013). DAF-16 targets are broadly classified into two categories: class I and class II. Class I DAF-16 targets are upregulated in the long-lived insulin/IGF-1-like signaling mutant *daf-2(-)* strain, while class II DAF-16 targets are downregulated in these animals (Tepper *et al*. 2013). Among the 1,784 genes found to be upregulated in either *hsb-1(-)* or *hsf-1* overexpression strains (Figure 5B), 293 genes, i.e., ∼16.4% were class I DAF-16 targets (Figure 7A) (Tepper *et al*. 2013). On the other hand, among the 164 genes identified as downregulated in either *hsb-1(-)* or *hsf-1* overexpression strains (Figure 5C), 29 genes, i.e., ∼17.7% were class II DAF-16 targets (Figure 7B) (Tepper *et al*. 2013). Interestingly, the proportion of class I DAF-16 targets in *hsf-1* overexpression-specific upregulated genes (256 out of 1,414 genes, i.e., ∼18.1%) was significantly higher than that expected by chance (*p* < 1.4 × 10^-19^ in hypergeometric test) (Figure 7A). In addition, the proportion of class II DAF-16 targets in *hsb-1(-)*-specific downregulated genes (23 out of 88 genes, i.e., ∼26.1%) was also significantly higher than that expected by chance alone (*p* < 5.45 × 10^-5^ in hypergeometric test) (Figure 7B). The number of DAF-16 target genes in the other categories (*hsf-1*-overexpression specific downregulated genes, *hsb-1(-)*-specific upregulated genes and genes upregulated or downregulated in both strains) were not found to be significantly higher than that expected by chance (*p* > 0.05 in hypergeometric tests) (Figures 7A and 7B).

**Figure 7.**
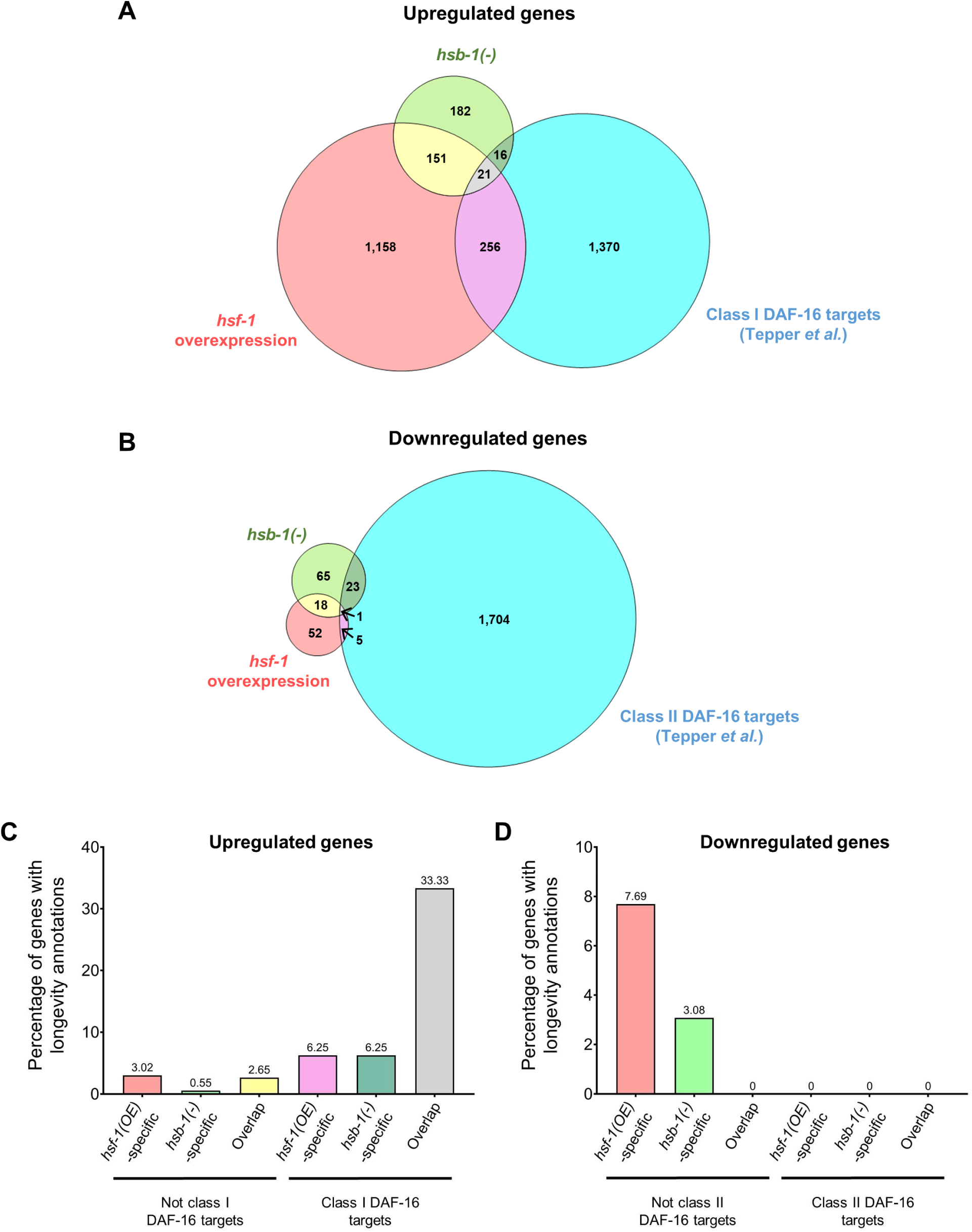
DAF-16 targets that are upregulated in both *hsb-1(-)* and *hsf-1* overexpression strains have a high proportion of known longevity genes. **(A)** Venn diagram showing overlap of genes upregulated in *hsb-1(-)* and *hsf-1* overexpression strains with previously reported class I DAF-16 targets (Tepper *et al*. 2013). **(B)** Venn diagram showing overlap of genes downregulated in *hsb-1(-)* and *hsf-1* overexpression strains with previously reported class II DAF-16 targets (Tepper *et al*. 2013). **(C, D)** Percentage of genes with known longevity functions among genes upregulated or downregulated in *hsb-1(-)* and *hsf-1* overexpression strains.

In a parallel approach, we determined the proportion of known longevity genes among the genes differentially expressed via HSB-1 inhibition or HSF-1 overexpression. In total, 64 out of 1,784 (∼3.6%) upregulated genes and 6 out of 164 (∼3.7%) downregulated genes were found to be annotated with longevity-related phenotypes on Wormbase (https://www.wormbase.org) (Figures 7A–D). The proportion of known longevity genes was significantly higher (24 out of 293, ∼8.2%) among class I DAF-16 targets identified in this study (*p* < 2.8 × 10^-5^in hypergeometric test) (Figures 7A and 7C). However, the higher proportion of longevity genes among DAF-16 targets might be due to confounding factors such as more extensive characterization of insulin/IGF-1-like signaling-mediated longevity in *C*. *elegans* compared to other less-studied life span regulating pathways. Nonetheless, even if the higher prevalence of known longevity genes among class I DAF-16 targets was taken into consideration, the percentage of genes previously implicated in life span regulation was still significantly higher among class I targets that were upregulated in both *hsb-1(-)* and *hsf-1* overexpression strains (7 out of 21 genes, i.e., ∼33.3%, *p* < 0.001 in hypergeometric test) (Figures 7A and 7C). In contrast, *hsf-1* overexpression-specific or *hsb-1(-)*-specific class I DAF-16 targets did not have a significantly higher proportion of known longevity genes (16 out of 256 genes, i.e., 6.25% and 1 out of 16 genes, i.e., 6.25%, respectively, *p* > 0.05 for both in hypergeometric tests) (Figures 7A and 7C). The class I DAF-16 targets that were upregulated via both HSB-1 inhibition and HSF-1 overexpression included *mtl-1, dod-3, dod-6, lys-7, asah-1* and *T20G5*.*8* (Tables S1 and S2), genes that are at least partially required for the life span extension phenotype in *daf-2* mutant animals (Murphy *et al*. 2003). However, not all known longevity genes identified in our transcriptomic analysis were DAF-16 targets. For example, the FOXA transcription factor *pha-4* is upregulated in both *hsb-1(-)* and *hsf-1* overexpressing animals (Tables S1 and S2), but is not required for the long life span of *daf-2* mutant animals (Panowski *et al*. 2007). Instead, *pha-4* has been shown to have an essential role in dietary restriction-induced longevity in *C*. *elegans* (Panowski *et al*. 2007). This suggests that HSF-1 not only influences the expression of longevity-promoting genes that act downstream of insulin/IGF-1like signaling, but also of genes involved in other life span-regulating pathways. Surprisingly, none of the class II DAF-16 targets identified in our study have been annotated with longevity-related phenotypes (Figures 7B and 7D). This indicates that in the context of life span regulation, DAF-16 and HSF-1 might suppress gene expression in animals via potentially distinct mechanisms (Tepper *et al*. 2013).

### Genes differentially expressed due to both HSB-1 inhibition and HSF-1 overexpression show a strongly correlated expression pattern

We speculated that genes that are differentially expressed in the same direction by both HSB-1 inhibition and HSF-1 overexpression might be contributing to their shared phenotype of HSF-1-associated life span extension in *C*. *elegans*. We visualized the fold-change in expression of the 214 genes that were differentially expressed in both *hsb-1(-)* and *hsf-1* overexpression strains relative to wild-type animals (Figures 5A and 8A). The heatmap showed that most of these genes have altered expression in the same direction with respect to the wild-type levels (Figure 8A; top hits are listed in Table 1). However, use of the same N2 control dataset for normalization could lead to a spurious correlation between the gene expression patterns in the two strains due to hidden covariates (Curran-Everett 2013). Hence, to avoid false-positive (type I error) similarities in expression patterns of the two strains, we re-calculated the direction of change in gene expression in the *hsb-1(-)* and *hsf-1* overexpression strains by using two independent biological replicates of N2 for normalization. Using this method, we found that 191 out of the 214 genes were either significantly upregulated (172 genes) or significantly downregulated (19 genes) in both *hsb-1(-)* and *hsf-1* overexpression strains relative to wild-type (Figure 8B). In contrast, less than 11 percent (23 out of 214) of these genes were found to be differentially expressed in opposite directions, i.e., upregulated in one strain but downregulated in the other (Figure 8B). The proportion of genes (89%) differentially expressed in the same direction in the two long-lived strains relative to wild-type was much higher than that expected by chance alone (chi-square value = 350.7, *p* < 0.0001), indicating the presence of common regulatory mechanisms for these genes in *hsb-1(-)* and *hsf-1* overexpressing animals.

**TABLE 1.**
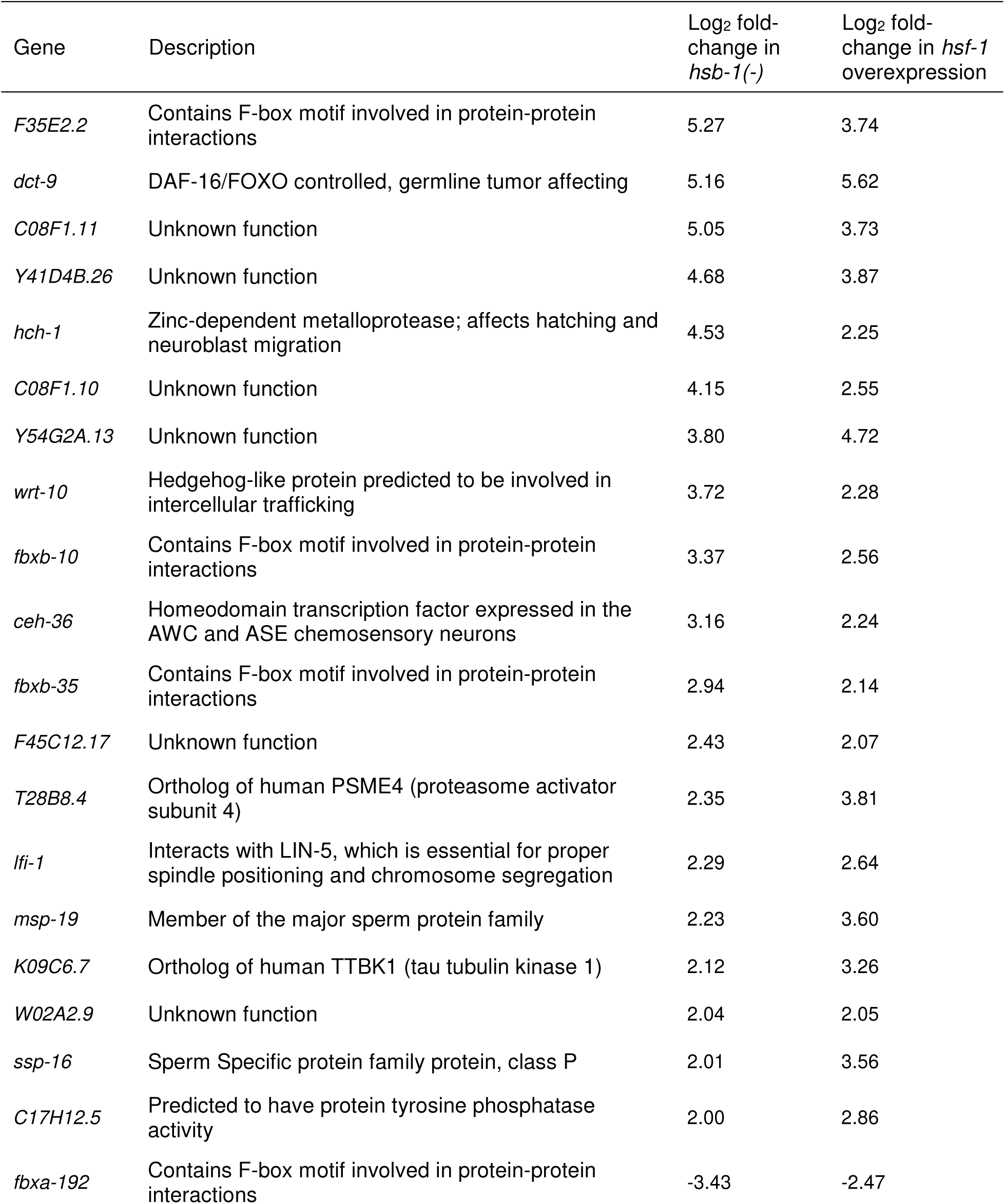
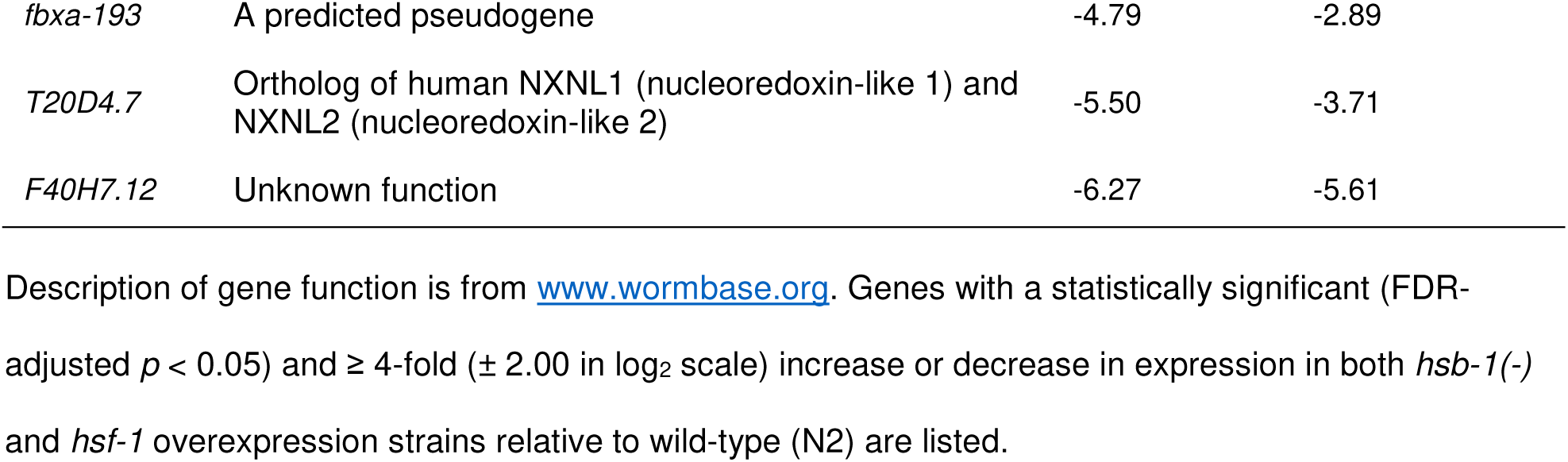
Top RNA-Seq hits that are differentially expressed in both *hsb-1(-)* and *hsf-1* overexpression strains relative to wild-type.

| Gene | Description | Log<sub>2</sub> fold-change in *hsb-1(-)* | Log<sub>2</sub> fold-change in *hsf-1* overexpression |
| --- | --- | --- | --- |
| *F35E2.2* | Contains F-box motif involved in protein-protein interactions | 5.27 | 3.74 |
| *dct-9* | DAF-16/FOXO controlled, germline tumor affecting | 5.16 | 5.62 |
| *C08F1.11* | Unknown function | 5.05 | 3.73 |
| *Y41D4B.26* | Unknown function | 4.68 | 3.87 |
| *hch-1* | Zinc-dependent metalloprotease; affects hatching and neuroblast migration | 4.53 | 2.25 |
| *C08F1.10* | Unknown function | 4.15 | 2.55 |
| *Y54G2A.13* | Unknown function | 3.80 | 4.72 |
| *wrt-10* | Hedgehog-like protein predicted to be involved in intercellular trafficking | 3.72 | 2.28 |
| *fbxb-10* | Contains F-box motif involved in protein-protein interactions | 3.37 | 2.56 |
| *ceh-36* | Homeodomain transcription factor expressed in the AWC and ASE chemosensory neurons | 3.16 | 2.24 |
| *fbxb-35* | Contains F-box motif involved in protein-protein interactions | 2.94 | 2.14 |
| *F45C12.17* | Unknown function | 2.43 | 2.07 |
| *T28B8.4* | Ortholog of human PSME4 (proteasome activator subunit 4) | 2.35 | 3.81 |
| *lfi-1* | Interacts with LIN-5, which is essential for proper spindle positioning and chromosome segregation | 2.29 | 2.64 |
| *msp-19* | Member of the major sperm protein family | 2.23 | 3.60 |
| *K09C6.7* | Ortholog of human TTBK1 (tau tubulin kinase 1) | 2.12 | 3.26 |
| *W02A2.9* | Unknown function | 2.04 | 2.05 |
| *ssp-16* | Sperm Specific protein family protein, class P | 2.01 | 3.56 |
| *C17H12.5* | Predicted to have protein tyrosine phosphatase activity | 2.00 | 2.86 |
| *fbxa-192* | Contains F-box motif involved in protein-protein interactions | -3.43 | -2.47 |
| *fbxa-193* | A predicted pseudogene | -4.79 | -2.89 |
| *T20D4.7* | Ortholog of human NXNL1 (nucleoredoxin-like 1) and NXNL2 (nucleoredoxin-like 2) | -5.50 | -3.71 |
| *F40H7.12* | Unknown function | -6.27 | -5.61 |

2 Description of gene function is from www.wormbase.org. Genes with a statistically significant (FDR-adjusted $p < 0.05$ ) and $\geq 4$ -fold ( $\pm 2.00$ in $\log_2$ scale) increase or decrease in expression in both *hsb-1(-)* and *hsf-1* overexpression strains relative to wild-type (N2) are listed.

**Figure 8.**
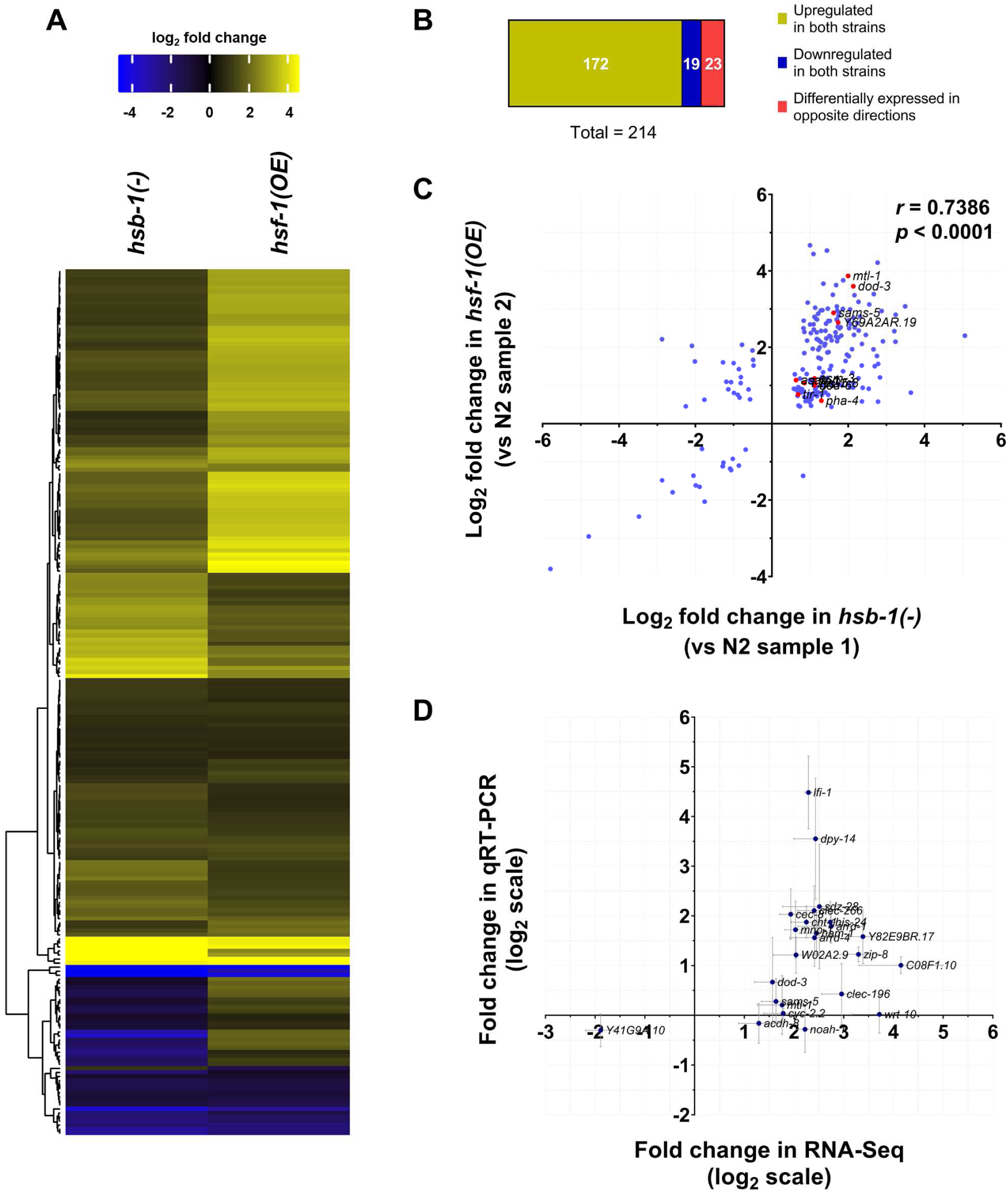
Correlation between expression pattern of genes that are differentially expressed in both *hsb-1(-)* and *hsf-1* overexpression strains. **(A)** Heatmap for fold-change in expression of 214 genes that are differentially expressed (fold-change ≥ 1.5) in both *hsb-1(-)* and *hsf-1* overexpression strains relative to wild-type (N2). Genes (rows) are clustered based on similarity in expression pattern. Colors show magnitude of change in gene expression (*yellow*: high, *black*: no change, *blue*: low). **(B)** Categorization of 214 genes shown in **(A)**. The three categories indicate genes that are upregulated in both *hsb-1(-)* and *hsf-1* overexpression strains, downregulated in both the strains or differentially expressed in opposite directions in the two strains. **(C)** Correlation between fold-change in expression pattern of 214 genes shown in **(A)**. Fold-change in gene expression for *hsb-1(-)* and *hsf-1* overexpression strains were calculated using two independent replicates of the wild-type strain for normalization. Red dots represent known longevity-regulating genes in *C*. *elegans*. **(D)** Fold-change in gene expression in *hsb-1(-)* animals relative to wild-type determined using RNA-Seq and quantitative RT-PCR. Expression data are shown for 24 transcripts that were among the top hits from RNA-Seq analysis. Data represent mean ± SEM for three biological replicates.

On comparison of the fold-change in expression of the 214 genes that were differentially expressed in both *hsb-1(-)* and *hsf-1* overexpressing strains, we found a strongly correlated expression pattern even when two independent wild-type control datasets were used for normalization (*r* = 0.7386, *p* < 0.0001) (Figure 8C). As described previously (Figure 8B), more than 89 percent of the 214 differentially expressed genes were found in the first and third quadrants of the scatter plot for relative change in gene expression (Figure 8C). Finally, we used quantitative RT-PCR to validate the magnitude of change in gene expression determined from RNA-Seq (Figure 8D). Among the 24 genes tested using quantitative RT-PCR, we found that most of the genes showed similar fold-change in expression in *hsb-1(-)* animals relative to wild-type as observed in our RNA-Seq data (Figure 8D). For some genes such as *C08F1*.*10, clec-196* and *lfi-1*, the magnitude of fold-change in expression (relative to wild-type) measured using quantitative RT-PCR did not precisely represent the value determined from RNA-Seq (Figure 8D). This is possibly due to the difference in transcript-binding sites and amplification efficiencies of the primers used for quantitative RT-PCR validation. However, for 20 out of the 24 genes tested, the change in direction of gene expression in the *hsb-1(-)* strain was the same irrespective of whether RNA-Seq or quantitative RT-PCR was used to measure transcript abundance (Figure 8D). Hence, our RNA-Seq analysis provides a fairly robust estimate of the transcriptome-wide changes in these long-lived *C*. *elegans* strains.

## DISCUSSION

Previous studies have established HSB-1 as an evolutionarily conserved negative regulator of HSF-1 that physically binds to the HSF-1 protein and sequesters it in an inhibitory complex (Satyal *et al*. 1998; Chiang *et al*. 2012). In this study, our findings collectively indicate that in the absence of HSB-1, transcriptional changes induced via activation of the HSF-1 protein result in life span extension in animals (Figure 9). We report increased *in vitro* binding activity of HSF-1 to its genomic target sequence in both *hsb-1(-)* and *hsf-1* overexpression strains (Figure 2C), but interestingly, HSF-1 protein level is not elevated in *hsb-1(-)* animals (Figure 2B). The higher *in vitro* HSE binding induced via HSF-1 overexpression is presumably due to the elevated HSF-1 protein level in that strain (Figures 2B and 2C). However, the increased *in vitro* HSE binding phenotype observed in *hsb-1(-)* animals can possibly be a consequence of yet unknown factors such as increased nuclear localization of HSF-1, acquisition of PTMs that favor DNA binding, or a combination of both (Figure 9). This concurs with the diverse HSR-independent roles of the HSF-1 transcription factor in different biological contexts (Li *et al*. 2017; Gomez-Pastor *et al*. 2018). Here, we identified the HSF-1-regulated transcriptome that is relevant for the longevity-promoting effects of this transcription factor in *C*. *elegans*. Though life span extension has been previously achieved via overexpression of HSF-1 in *C*. *elegans* (Hsu *et al*. 2003; Morley and Morimoto 2004; Baird *et al*. 2014), our findings suggest that the beneficial effects of HSF-1 activation upon longevity could also be achieved via modulating the expression of a limited number of its transcriptional targets (Figures 4A–D).

**Figure 9.**
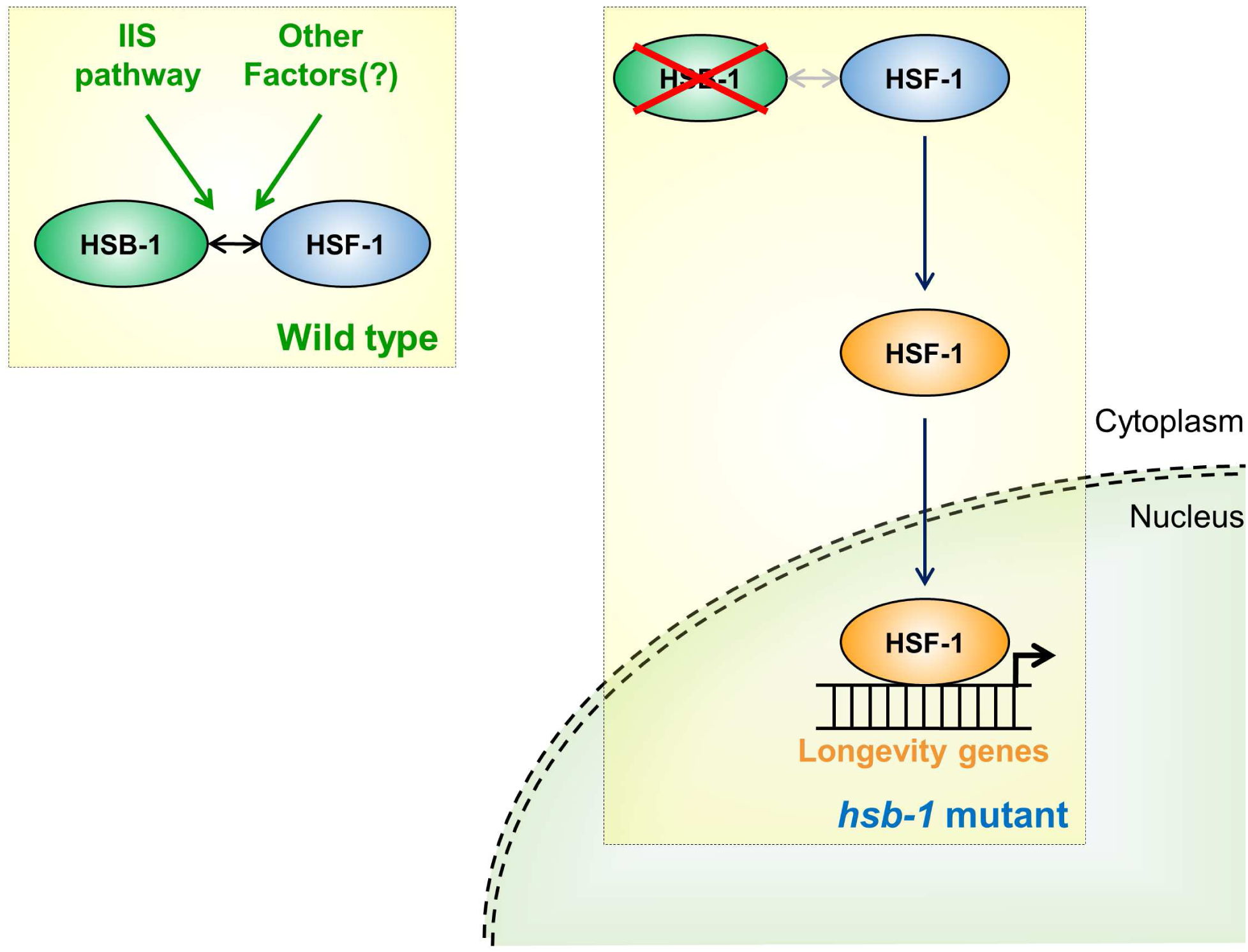
Model for longevity due to an HSF-1-induced altered transcriptional profile in the absence of HSB-1 regulation. The proposed model indicates HSF-1-associated regulation of longevity in *hsb-1(-)* mutant animals. In wild-type animals, HSB-1 binds to HSF-1 and limits its transcriptional activation potential. As shown previously (Chiang *et al*. 2012), this binding is promoted by insulin/IGF-1-like signaling (IIS) pathway and potentially, also by other unknown factors. In the absence of HSB-1 regulation in *hsb-1(-)* mutant animals, HSF-1 protein is activated (blue to orange), translocates to the nucleus and promotes expression of its target genes. This HSF-1-induced change in the transcriptional profile results in increased life span of *hsb-1(-)* animals.

Our transcriptomic analyses show that a sizeable fraction of differentially expressed genes in the *hsb-1(-)* mutant and *hsf-1* overexpression strains have a strongly correlated expression pattern (Figures 8A and 8C), suggesting similar transcriptional regulation of a subset of HSF-1 target genes in both these strains. Incidentally, though the heat shock response genes *hsp-16*.*2* and *hsp-70* showed roughly two-fold increase in expression compared to wild-type in both *hsb-1(-)* and *hsf-1* overexpression strains in our quantitative RT-PCR experiments (Figures 2D and 2E), these *hsp* genes were not identified to be differentially expressed in our transcriptome-wide RNA-Seq comparison (Tables S1 and S2). We found that this was due to very low sequencing read counts for the *hsp* genes across all the samples. Quantitative RT-PCR data indicated that the expression levels of these *hsp* genes were ∼10,000-fold lower than that of housekeeping genes in non-stressed conditions (data not shown). Hence, higher sequencing depth can potentially identify the difference in transcript levels of *C*. *elegans* genes that have extremely low basal expression in normal physiological conditions. Among the genes identified in our transcriptomic analysis, it remains to be determined which of those are causally linked to the life span extension phenotype in the *hsb-1(-)* mutant. It will be informative to estimate the extent to which inactivation of individual genes affects the life span of *hsb-1(-)* animals. However, the longevity phenotype associated with HSB-1 inhibition is most likely an additive effect of the contributions of numerous HSF-1 target genes identified in this study and might also involve epistatic interactions between those downstream targets. Moreover, it is possible that certain differentially expressed transcripts in the *hsb-1(-)* strain are regulated in an HSF-1-independent manner. Characterization of how HSF-1 genomic occupancy is altered in *hsb-1(-)* animals will help to identify loci that are directly under the control of HSB-1/HSF-1 pathway.

Nonetheless, our RNA-Seq analysis has identified several known longevity-promoting genes, such as *mtl-1, dod-3, lys-7* and *pha-4*, that are upregulated in the HSF-1-dependent long-lived strains (Figure 8C, and Tables S1 and S2). Most of these longevity genes act downstream of the insulin/IGF-1-like signaling pathway in *C*. *elegans* (Murphy *et al*. 2003; Tepper *et al*. 2013), while *pha-4* is involved in regulation of dietary restriction-induced macroautophagy (Hansen *et al*. 2008). In addition, our transcriptome-wide comparison also revealed increased expression of more than 1,500 *C*. *elegans* genes in *hsf-1* overexpressing animals (Figures 4B and 5B). Since the number of upregulated genes was comparatively much lower in the *hsb-1(-)* mutant strain (Figures 4A and 5B), we conclude that HSF-1-induced longevity can be achieved via altered expression of a much smaller subset of HSF-1 target genes. A recent transcriptome-wide study has reported that HSF-1 regulates the expression of more than 2,000 genes independently of the heat shock response in *C*. *elegans* (Brunquell *et al*. 2016). Many of these genes are involved in maintenance of nematode cuticle structure and other metabolic processes such as mitochondrial function, regulation of growth etc. Interestingly, roughly 90% of these HSF-1-regulated transcripts do not overlap with a gene expression dataset for aging-induced transcriptional changes in *C*. *elegans* (Brunquell *et al*. 2016). This further indicates that ectopic HSF-1 activation might inadvertently affect numerous biological pathways in addition to the ones involved in longevity regulation. Since mammalian HSF1 has been shown to support progression of highly malignant cancers (Dai *et al*. 2007; Mendillo *et al*. 2012), it is crucial to develop strategies that can selectively modulate the activity of this transcription factor to promote longevity in animals without producing undesired outcomes.

In summary, our study provides a better understanding of the function and regulation of heat shock factor in non-stressed physiological conditions. The findings from this study can potentially guide the design of targeted therapies to exploit the untapped potential of this evolutionarily conserved longevity-promoting factor in delaying age-associated physiological decline in complex organisms.

## ACKNOWLEDGEMENTS

We thank Dr. Shweta Ramdas for her helpful comments on the manuscript. Some *C*. *elegans* strains were obtained from the *Caenorhabditis* Genetics Center (University of Minnesota), which is supported by the NIH Office of Research Infrastructure Programs (P40 OD010440). The following funding sources supported this research: National Institute of Aging (R01 AG028516) and Ministry of Science and Technology of Taiwan (107-2628-B-010-002) (to A-L. H.).

